# Color and gloss constancy under diverse lighting environments

**DOI:** 10.1101/2022.12.09.519756

**Authors:** Takuma Morimoto, Arash Akbarinia, Katherine Storrs, Jacob R. Cheeseman, Hannah E. Smithson, Karl R. Gegenfurtner, Roland W. Fleming

**Affiliations:** Department of Psychology, Justus-Liebig-Universität Gießen, Germany; Department of Experimental Psychology, University of Oxford, Oxford, UK; School of Psychology, University of Auckland, New Zealand

**Keywords:** color, gloss, perceptual constancy, directional lighting environments, asymmetric matching

## Abstract

When we look at an object, we simultaneously see how glossy or matte it is, how light or dark, and what color. Yet, at each point on the object’s surface, both diffuse and specular reflections are mixed in different proportions, resulting in substantial spatial chromatic and luminance variations. To further complicate matters, this pattern changes radically when the object is viewed under different lighting conditions. The purpose of this study was to simultaneously measure our ability to judge color and gloss using an image set capturing diverse object and illuminant properties. Participants adjusted the hue, lightness, chroma, and specular reflectance of a reference object so that it appeared to be made of the same material as a test object. Critically, the two objects were presented under different lighting environments. We found that hue matches were highly accurate, except for under a chromatically atypical illuminant. Chroma and lightness constancy were generally poor, but these failures correlated well with simple image statistics. Gloss constancy was particularly poor, and these failures were only partially explained by reflection contrast. Importantly, across all measures, participants were highly consistent with one another in their deviations from constancy. Although color and gloss constancy hold well in simple conditions, the variety of lighting and shape in the real world presents significant challenges to our visual system’s ability to judge intrinsic material properties.

## 1. Introduction

In our everyday lives, we often identify and interact with objects across major changes in lighting—for example, returning at sunset to a car we parked in the morning, or carrying a familiar coffee mug from the kitchen to the balcony. In such circumstances, we are not usually struck by the impression that the color of the car’s paint, or the gloss of the mug’s glaze, have changed. Yet judging the color and gloss of a surface—especially across different lighting conditions—poses hard computational challenges. Firstly, these perceptual quantities result from two physically distinct aspects of the surface’s reflectance, which are confounded in the retinal image. Surface color typically depends primarily on diffuse reflection in which the incident illumination is modified through interaction with pigments, whilst gloss stems from specular reflection, which is a direct reflection of the illuminant. At each point on the object’s surface, diffuse and specular reflections are mixed in different proportions, resulting in substantial spatial chromatic and luminance variations. In the demonstration in Figure 1 (a), pixel colors dramatically vary among three selected points on the surface of the object, showing that there is no simple mapping between the cone excitations at a single pixel and the color of the object. Furthermore, when we created this image (using computer graphics rendering techniques), we placed an object under a specific lighting environment, applied specific diffuse and specular reflectance properties to the object, and fixed the camera at a specific viewpoint (panel (a) upper-right). Changing any of these underlying scene parameters would alter the color distribution across the same object’s surface in the resultant image. As shown in Figure 1 (b), pixel colors of the same three surface points change substantially when the same object is viewed in the same pose, but in different lighting environments. The reader may also have the impression that the color and glossiness of the object appear somewhat different, even though the material properties are identical across the three objects.

**Figure 1:**
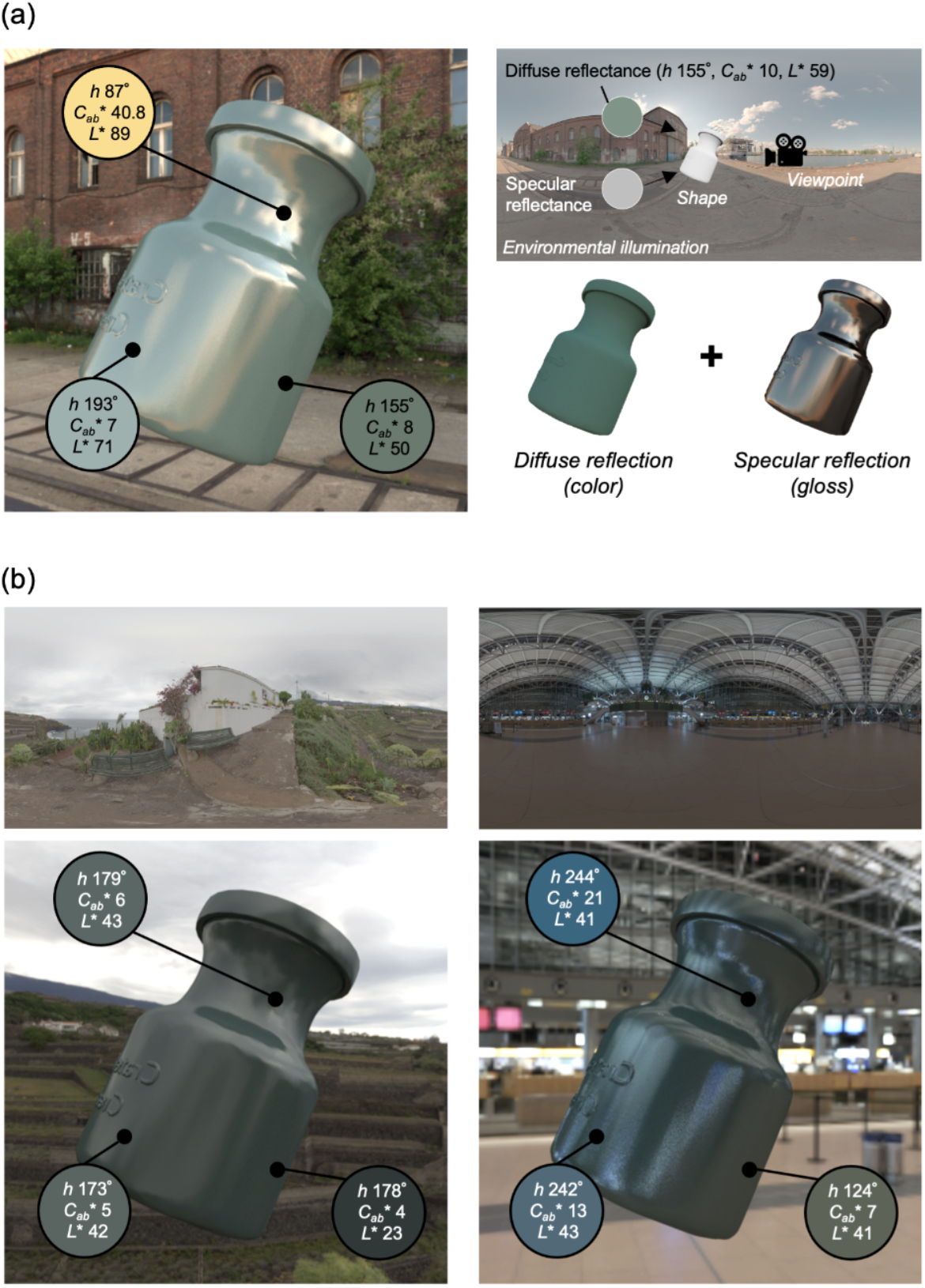
Graphical illustration of challenges to judging the color and gloss of a three-dimensional object placed under a complex lighting environment. (a) As shown to the upper-right, we placed an object under an environmental illumination, applied a diffuse reflectance which has a fixed hue, chroma and lightness (denoted as *h, C*_*ab*_* and *L*\*, respectively) and a specular reflectance, set a view-point and rendered the object image using computer graphics techniques. The rendered image is shown on the left side. Hue, chroma and lightness vary largely across the three selected pixels at different regions on the object’s surface. (b) Effects of lighting environments (overcast environment on the left and indoor environment on the right) on the color variation at the same three locations. Note that the objects in panels (a) and (b) have identical material properties though their appearance may differ across lighting environments.

Color constancy has been a core domain of human color vision research (Foster, 2011; Smithson, 2005; Hurlbert, 2007). Many of the early studies used simplistic visual stimuli that were flat, matte, and uniformly illuminated, but more recent studies have begun to use more complex stimuli that better represent natural visual environments (Brainard & Maloney, 2004; Witzel & Gegenfurtner, 2018; Mizokami 2019). One common research question is whether the use of real three-dimensional objects increases the degree of color constancy, compared to flat and uniformly-colored surfaces or images presented on a computer monitor (Hedrich, Bloj & Ruppertsberg 2009; de Almeida, Fiadeiro & Nascimento, 2010; Morimoto et al. 2017). Color and lightness constancy in the presence of specular reflection have also been well-studied, and several studies have found that the presence of specular reflections can improve color constancy (Hurlbert et al. 1991; Yang & Maloney, 2001; Yang & Shevell, 2003; Snyder, Doerschner & Maloney, 2005; Granzier, Vergne & Gegenfurtner 2014; Lee & Smithson, 2016; Nagai, et al. 2017) while other works found the effect only in limited conditions (Xiao et al. 2012; Wedge-Roberts et al. 2020).

The effect of object and lighting properties on perceived gloss has also been the subject of several studies, although the term “constancy” is not always explicitly stated. Many factors that can alter perceived gloss have been identified: e.g., object motion (Doerschner et al., 2011), body color (Wendt et al., 2010), surface curvature (Ho, Landy, & Maloney, 2008), and lighting environment statistics (Fleming, Dror & Adelson, 2003; Obein, Knoblauch, & Viéot, 2004; Pont & te Pas, 2006; Adams et al., 2018). Furthermore, systematic perturbation of lighting and material properties have been used to quantify their influences on perceived gloss level (Zhang, Ridder, & Pont, 2018; Zhang et al. 2019; Zhang et al., 2020). Motoyoshi and Matoba (2012) used scenes containing a statue along with other objects. They showed failures of gloss constancy and that surrounding context information had virtually no effect on matching results, implying that participants do not use contextual information to discount the influence of illumination.

In addition to these constancy perspectives, specific visual computations that may underlie gloss perception have been proposed in recent decades (Landy, 2007; Chadwick & Kentridge, 2015). One primary discussion point is whether the visual system makes use of summary statistics extracted from a given image, rather than reconstructing the entire optical input (Fleming, 2014; Fleming, 2017; Nishida, 2019). Candidate cues to gloss include skewness of the luminance histogram (Motoyoshi et al., 2007; Sharan, et al. 2008; Anderson & Kim, 2009; Kim & Anderson, 2010), its standard deviation (Wiebel, Toscani & Gegenfurtner, 2015), the magnitude of luminance gradients in an image (Sawayama & Nishida, 2018), and more complex image metrics computed from specular reflection patterns (Marlow, Kim & Anderson 2012; Marlow & Anderson, 2013). Perceived gloss is non-linearly related to underlying physical quantities such as the proportion of light reflected in a specular manner, and the discriminability of gloss as a function of specular reflectance has been recently investigated (Cheeseman et al. 2021).

Despite these extensive investigations into color constancy and gloss constancy as separate domains, very few studies have simultaneously measured color *and* gloss constancy (lightness and gloss, Olkkonen & Brainard, 2010, Hansmann-Roth & Mamassian, 2017; gloss and hue variation between green and blue, Radonjić et al. 2018, Brainard, Cottaris & Radonjić 2018, Radonjić et al. 2019). No study has measured all dimensions of perceived color (hue, chroma and lightness) and gloss at the same time. Yet, in daily life, when we look at an object, both percepts naturally occur together, and we do not judge each separately unless explicitly asked to do so. The purpose of this study was to directly measure our ability to judge color and gloss using a synthetic image set produced using physically-based ray-tracing techniques from computer graphics that captures large variations of object properties and illuminant properties. For this, we used a well-established asymmetric matching task in which two images were presented side-by-side on a computer screen. Participants were asked to adjust the color and gloss of a right reference object so that it appears to be made of the same material as a left test object. The critical manipulation is that the two objects were presented under different lighting environments, and thus participants needed to take this difference into account to achieve accurate matching of physical reflectance parameters. This methodology has been extensively used in perceptual constancy studies and shown to be an effective constancy task (Arend & Reeves 1986; Brainard & Wandell, 1992). For instance, Olkkonen & Brainard (2010) jointly measured perceived lightness and gloss of smooth spheres placed under real-world lighting environments. They showed that participants accurately judged lightness, but perceived gloss varied substantially across lighting environments. There are a few other studies using the asymmetric matching task which did not consider the influence of illuminant changes, but which are nonetheless relevant to the present study. Xiao and Brainard (2008) showed that humans do not simply use the mean color across the whole object to determine the overall color impression of a three-dimensional glossy object. Strong interactions between specular reflection, chroma and lightness were found (Honson et al. 2020; Isherwood et al., 2021). Color matching using different types of materials (papers, sponges, wool, candles and porcelain) revealed that hue perception is highly stable, while chroma and lightness were influenced by material types (Giesel & Gegenfurtner 2010).

This study built upon these past efforts in two ways: (i) participants were asked to judge three color dimensions (hue, lightness and chroma) and gloss at the same time, (ii) we used a substantially wider variety of lighting environments and object shapes to capture the complex behavior of constancy mechanisms (including successes and errors) in a large stimulus space. Two psychophysical experiments were performed. The first experiment measured perceived color and gloss from single objects, under different lighting environments, where each object was randomly assigned a unique color and gloss level. The second experiment followed up the failure of gloss constancy observed in the first experiment by systematically exploring the effects of object shape and lighting environment on perceived gloss.

## 2. General Method

### 2.1 Apparatus

Both experiments were computer-controlled and all images were displayed on a 24-inch LCD monitor (ColorEdge CG2420, 1920×1200 pixels, frame rate 60 Hz; EIZO, Ishikawa, Japan) in 10 bits per color channel (red, green and blue). We performed gamma correction and spectral calibration using a spectroradiometer (CS-2000; KONICA MINOLTA, Inc., Tokyo, Japan). Experimental code was written in MATLAB using custom-built functions as well as functions provided in PsychToolbox-3 (Brainard, 1997).

### 2.2 Participants

Ten naive participants were recruited for Experiments 1 and 2. The ratio of female to male was 2.33 for Experiment 1 and 1.50 for Experiment 2. The age of participants ranged from 21 to 32; their mean ± 1.0 S.D. was 25.6 ± 2.83 for Experiment 1 and 24.0 ± 3.33 for Experiment 2. Four participants completed both experiments. The experiments were approved by a local ethics committee at Justus Liebig University Giessen in accordance with the Helsinki Declaration (6th revision, 2008). Before the experiments, all participants were screened for normal color vision using Ishihara pseudoisochromatic testing plates (Ishihara, 1973). All participants were undergraduate students at Justus Liebig University Giessen, Germany and paid for their time.

### 2.3 Task

We used an asymmetric matching task. As shown in Figure 2, a test image and a reference image were presented side by side, separated by 7.1° of visual angle. Participants were asked to adjust the color and gloss of the right reference object until it appeared to be made of the same material as the left test object. The adjustment of color and gloss were done in the “object” space rather than the “image” space. In other words, for the color setting, participants changed an underlying diffuse reflectance of the reference object by adjusting hue, lightness (*L*\*) and chroma (*C*\*_ab_) (as defined in the *L*\**a*b\** color space, for the reflectance under equal energy white) which updated the weights of reflectance basis functions (described below) to produce a composite reflectance with the desired color properties. For the gloss setting, participants changed an underlying specular reflectance by adjusting a parameter *c*, a perceptually linear gloss scale developed by Pellacini et al. (2000) (‘*Pellacini’s c*’ hereafter). We used these approximately perceptually linear scales for color and gloss adjustment because we predicted that participants might find it easier if increasing or decreasing a single step would have approximately equal perceptual effect at any point on each parameter range. Participants used eight buttons, each corresponding to increasing or decreasing values of one of the four parameters. A beep was provided when the value reached the limit of the prepared range for lightness, chroma, and *Pellacini’s c*. Hue is a circular variable, and thus there was no range limit. Participants binocularly viewed stimuli presented on a flat screen. During the matching task, participants were allowed to move their eyes freely between the test image and reference image. There was no time limit for each trial. In this study the right reference object served as a scale to quantify participants’ subjective experiences, and thus the shape and the lighting environment in the reference image stayed the same during the whole experiment, and only the left test image changed from one trial to the next. A key feature of this task was that the two objects were placed under different lighting environments. Thus participants needed to discount the effect of lighting on appearance to make accurate color and gloss matches. Pixel colors have a complex relationship to the physical parameters applied to the test object because they are influenced by the lighting environment, as shown in Figure 1. However, a perfectly color- and gloss-constant observer should be able to estimate physical parameters from the test image regardless of the lighting environment and assign the same parameters to the reference image. Details of test images and reference images will be explained in the subsequent section.

**Figure 2:**
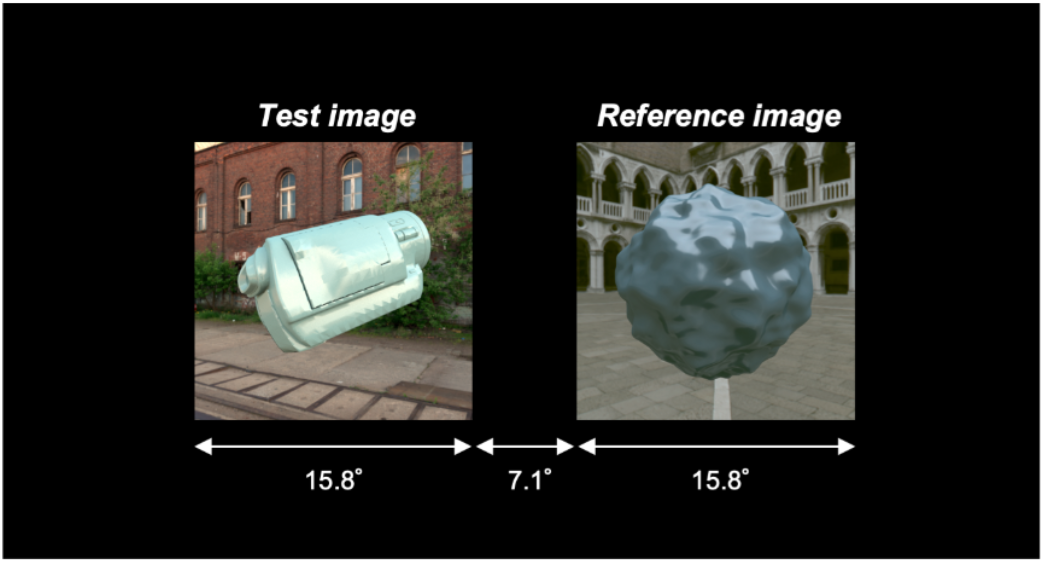
Stimulus configuration for asymmetric matching in this study. The task of participants was to adjust an underlying diffuse reflectance to change the color and an underlying specular reflectance to change the gloss of the reference object presented in the right reference image until it appears to be made of the same material as the test object presented in the left test image. Test images changed from one trial to another, but the reference image stayed the same throughout the experiment. White text and arrows were not presented during the actual experiment.

### 2.4 Experimental stimuli

#### 2.4.1. Rendering

All images used in this study were generated using the physically-based renderer Mitsuba (Jakob, 2010). A single image contained one floating object shape, to which we applied a diffuse reflectance and a specular reflectance using the Ward reflectance model (Ward, 1992). For the illumination we used image-based lighting (as shown in Figure 1), which we call ‘environmental illumination’ in this study, where each pixel in the environmental illumination map conveys information about a light ray that reaches a single point in the scene from a particular direction. The map thereby depicts light coming from every possible direction in the environment. Environmental illuminations used in this study were all originally RGB images, and Mitsuba was used to convert these to hyperspectral images during the rendering process (Smith, 2000). To maximize the accuracy of the rendering process, all images were spectrally rendered from 400 nm to 720 nm in 10 nm steps (31 spectral channels). All images were rendered in 512×512 pixel resolution. Rendered hyperspectral images were converted to the XYZ color space to calculate image statistics, and to the calibrated RGB color space such that displayed images reproduced the XYZ values from the rendered images.

#### 2.4.2 Reference images

Measuring perceived color and gloss using the asymmetric matching task required a reference scene in which participants can adjust the physical reflectance parameters of the reference object in a continuous fashion. To construct the reference scene, we selected (i) environmental illumination, (ii) shape of the reference object, and (iii) diffuse and specular reflectances as follows.

In selecting the reference environment, we wanted an environment in which chromatic variation is low and a sufficient amount of light directly hits the objects embedded in this environment producing highly visible specular reflections, which we expected could help participants to easily judge both color and gloss of the reference object. We also wanted to choose an environment in which the reference object appears highly glossy, as otherwise participants may not find a satisfactory match with a test object even at the highest gloss level allowed for the reference object. To establish this, we plotted diagnostic metrics, for publicly available maps, as shown in Figure 3. Panel (a) shows the selected environmental illumination map (‘Uffizi Gallery, Italy’) downloaded from a publicly available database (Debevec, 1998; https://vgl.ict.usc.edu/Data/HighResProbes/, accessed on 15th March 2022) along with statistical characterization of the illumination. The upper-right plot shows a luminance histogram of all pixels together with some basic statistics: mean, standard deviation (s.d.), skewness (skew.), kurtosis and Xia’s diffuseness metric (Xia. et al. 2017) which quantifies how much the lighting environment is directionally uniform from 0 (point light source) to 1 (fully uniform). This lighting environment has low diffuseness because upper and lower hemifields have largely different directional lighting patterns. The lower-left plot shows 10% pixel distribution on *a*\**b*\* chromatic plane where equal energy white is set to the origin. The lower-right panel shows a power spectrum, analyzed by spherical harmonic decomposition, with a negative slope of the regression line about -2 that was shown to be typical for outdoor environmental illuminations (Ron et al. 2004).

**Figure 3:**
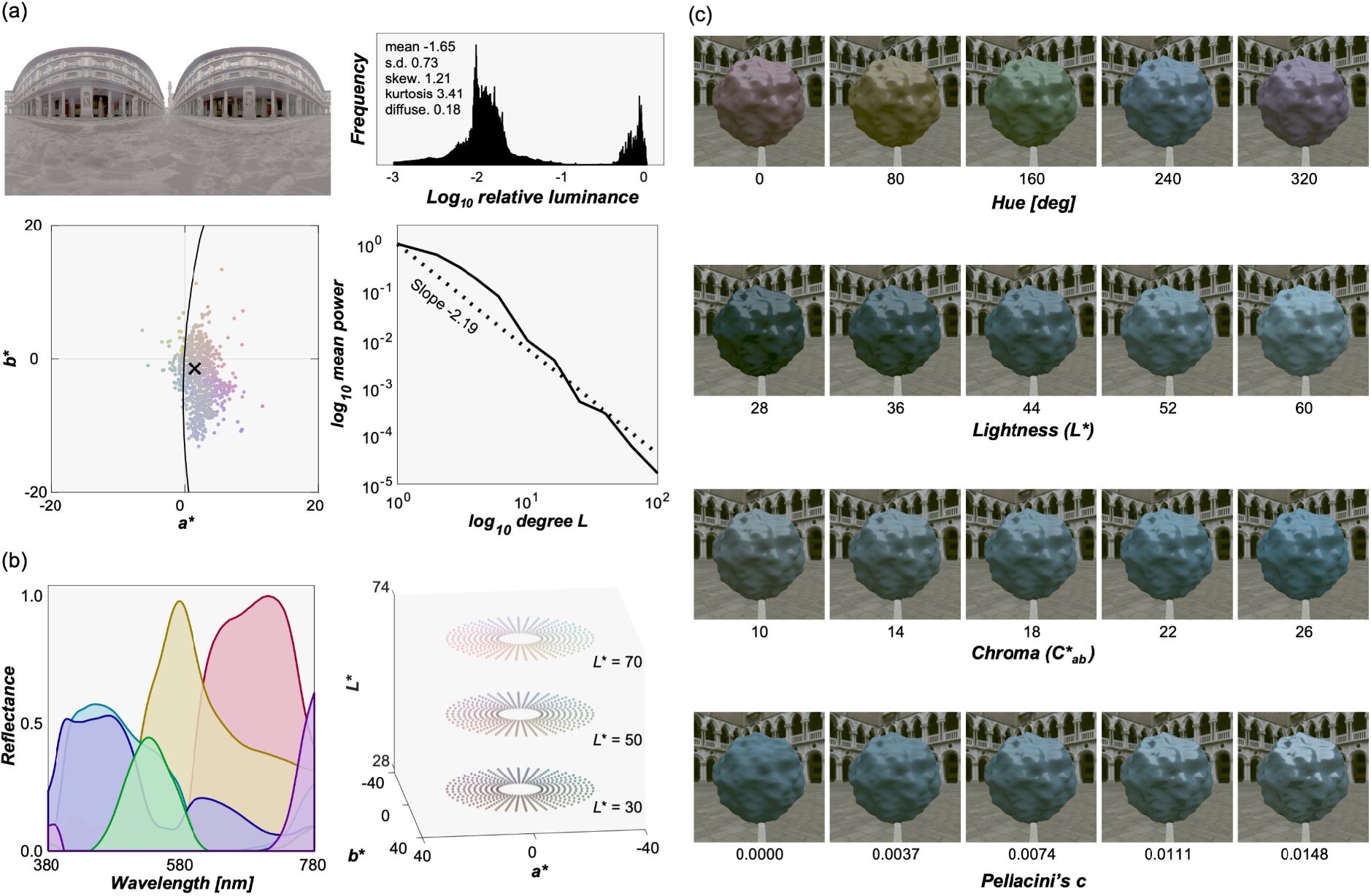
(a) Reference environmental illumination used for the reference image and its statistical characterization. The upper right plot shows the luminance histogram of all pixels, along with some image statistics (mean, standard deviation, skewness, kurtosis and Xia’s diffuseness metric). The lower left plot shows the *a*\**b*\* chromatic distribution of 10% pixel colors that were randomly sampled. The lower right plot shows the power spectrum analyzed by spherical harmonic decomposition. (b) Six basis reflectance functions extracted from 1,269 matte Munsell color chips using non-negative matrix factorization. Subset of surface colors assigned to the object in the reference images at example lightness planes (*L*\* = 30, 50 and 70). There were in total 21,600 colors, allowing participants to explore the stimulus space using method of adjustment. (c) Some example reference images, drawn from the four dimensional stimulus space of color and gloss adjustments.

For the reference shape, we chose a simple bumpy sphere whose surface roughness was fixed at 0.05 as defined in Mitsuba. We chose this shape as a standard shape again because some degree of bumpiness increases the spatial contrast of specular reflection between concave and convex regions, helping participants to detect where the specular reflections are, and high spatial-frequency geometries have been associated with better material recognition accuracy (Lagunas et al., 2021).

For a continuous adjustment of color and gloss, we needed to systematically control the underlying physical reflectances, separately for diffuse reflectance and specular reflectance. Our approach was to separately render diffuse and specular images which were linearly summed to produce a colored glossy object, which provided a good enough approximation for the geometry considered here.

We first generated a specular image that has only a specular reflection component by setting the specular reflectance to 1.0 and diffuse reflectance to zero across all wavelengths. To modulate the gloss level of test images, this specular image was multiplicatively scaled up and down by a single factor. As already described, participants adjusted the gloss level by changing Pellacini’s c, and this parameter was directly converted to specularity *p*_*s*_, a parameter defined in the Ward reflectance model. The scalar quantity *p*_*s*_ was used to scale the specular image at each wavelength. We sampled Pellacini’s c from 0 (matte) to 0.149 (highly glossy), which corresponded to 0 and 0.0999 in specularity *p*_*s*_, respectively. Pellacini’s c captures the lightness-dependent nature of perceived gloss and thus requires a lightness value for the conversion. For this, we used a value of 50, which is roughly equivalent to the mid value of the prepared lightness range (detailed below).

Our approach to define diffuse reflectances was to first specify colors in terms of hue, lightness and chroma in *L*\**a*\**b*\* color space, and find a spectral reflectance, by combining basis reflectance functions extracted from the Munsell color system, that produces the desired hue, lightness and chroma when placed under an equal energy white light (X = Y = Z = 100).

We first applied non-negative matrix factorization to 1,269 matte Munsell color chips (https://sites.uef.fi/spectral/munsell-colors-matt-spectrofotometer-measured/, accessed on 15th March 2022) to obtain 6 basis spectral reflectance functions as shown in the left panel of Figure 3 (b). Then, we sampled 21,600 colors in *L*\**a*\**b*\* color space (subset is shown in the right plot of Figure 3 (b)): 90 hue values from 0° to 356° in 4° steps; 24 lightness values from 28 to 74 in 2 steps; 10 chroma values from 8 to 26 in 2 steps. Then, for each color, we searched for the optimal weights for 6 basis reflectances so that the weighted summation of the basis functions produced the desired *L*\**a*\*b* values under equal energy white light (X = Y = Z = 100). For the optimization we set a condition that the resultant reflectance value should be between zero and one at any wavelength to meet physical constraints. This process generated 21,600 spectral diffuse reflectances, and using these we rendered 21,600 matte reference images. We used 36 of these reflectances to generate test images in Experiment 1 as described in the subsequent section. These diffuse images along with the aforementioned single specular image enabled continuous adjustment of the color and gloss of the objects in such a way as to achieve approximately uniform coverage of the perceptual space, at least as expected under reference viewing conditions. Figure 3 (c) shows some example reference images.

When we rendered scenes in the Mitsuba renderer, the object was placed at the center of the scene and environmental illumination was applied to the scene. Then, the camera was set at the same height as the object and looked directly at the object. Environmental illumination generally had high dynamic range, and had we selected the viewpoint (i.e. camera position) at random it would have often created an image in which the object region was too dark to see. Thus, we rendered a mirrored sphere from different viewpoints (0° to 330° in 30° steps) and picked the viewpoint at which the mean luminance over the sphere was the highest. We also made sure that there were no pixels with particularly intense lights in the surrounding non-object region of the image. Pixel values in the raw hyperspectral images from Mitsuba were arbitrarily defined because pixel values in the environmental illumination have no units. Thus, after converting the raw hyperspectral image to a linear monitor RGB image, we scaled the whole image by the 99th percentile pixel value across all reference images. We applied the same scaling value to all reference images to equate the light level across images. Because of these selection processes, no tone mapping was applied to any of the reference images. These scaled linear RGB images were gamma-corrected and used for the experiment.

#### 2.4.3 Test images

We first gathered environmental illuminations from multiple online databases (including Adams et al. 2016; Debevec, 1998; and https://hdrmaps.com/freebies/) from which we selected 12 lighting environments that cover a diverse variation of natural lighting. Figure 4 shows selected lighting environments along with the luminance histogram of all pixels and 10% pixel chromatic distributions on *a*\**b*\* plane for (a) 6 outdoor scenes including sunny days (left 4 scenes) and cloudy days (right 2 scenes) and (b) 6 indoor scenes.

**Figure 4:**
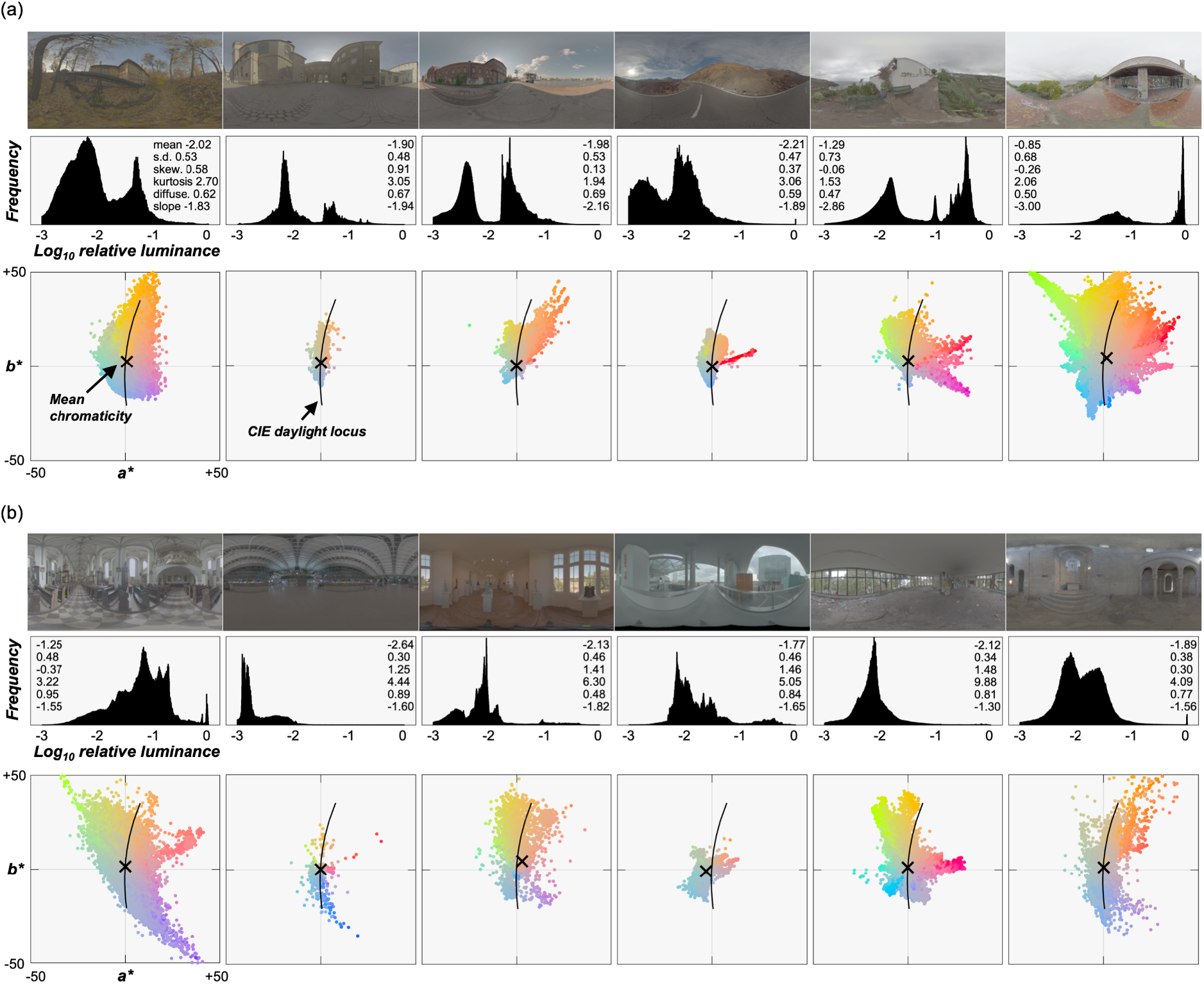
(a) 6 outdoor lighting environments (left 4 environments are sunny days and the right 2 environments are overcast days) and (b) 6 indoor lighting environments, together with the luminance histogram of all pixels and the *a*\**b*\* chromatic distribution of 10% of pixel colors sampled at random. The numbers in the histograms are mean, standard deviation (s.d.), skewness (skew.), kurtosis, Xia’s diffuseness metric (diffuse.) and slope of the power spectrum computed by spherical harmonic decomposition. Maximum luminance in each image was normalized to 1.0 (zero on a logarithmic scale) to allow the comparison across environments in this figure, but note that for actual test images each environment was scaled differently from this figure (detailed in main text). The intersection of horizontal and vertical thin gray lines denotes the white point (X=Y=Z=100) of the color space. The black cross symbol depicts mean *a*\**b*\* chromaticity across the plotted 10% of pixels. The black solid line shows the CIE daylight locus.

To expand the diversity of the selected lighting environments to unnatural domains, we applied two independent manipulations to each lighting environment. Firstly, we rotated the *a*\**b*\* chromatic distribution of the original lighting environment by +90° while keeping the *L*\* distribution the same, producing ‘gamut-rotated lighting environment’. Chromaticities in natural lighting environments tend to cluster along a blue-yellow axis known as the CIE daylight locus (Judd et al., 1964; Hernández-Andrés et al. 2001), and this manipulation made the distribution run along an orthogonal red-green axis. If our visual system uses a prior about a typical illuminant to achieve perceptual constancy (Delahunt & Brainard, 2004; Pearce et al. 2014; Weiss, Witzel & Gegenfurtner 2017), we may observe higher errors in participants’ settings under these chromatically atypical environments. After the rotation, some pixels in the map went outside the chromatic gamut of the experimental monitor and were thus likely to produce out-of-gamut pixels in the rendered images. For those pixels, the chroma value was decreased until the colors came inside the chromatic gamut. When selecting the original lighting environments we made sure that such pixels were always less than 5%. The second manipulation was to distort the directional structure of each lighting environment using a phase scrambling technique via spherical harmonic decomposition. This manipulation normally changes the color distribution of the image, but we kept the distribution of chromaticity and luminance by histogram matching the phase-scrambled environment maps to the originals. Again, if our visual system uses a prior about the directional structure, such as ‘light from above’ (e.g. Ramachandran, 1988; Morgenstern et al. 2011), we might observe poor perceptual constancy under these environments because intense lights could hit the object from every direction. The resultant 36 lighting environments (12 natural, 12 gamut-rotated, 12 phase-scrambled) were used in Experiment 1. Twelve natural lighting environments were used in Experiment 2.

For Experiment 1, for each of the 36 lighting environments, we placed in the scene a randomly-selected object from a dataset of 3-D meshes of everyday objects (purchased from Evermotion, https://evermotion.org/), and physical reflectance parameters (hue, chroma, lightness and Pellacini’s c), and selected the viewpoint in the same way we selected the viewpoint for the reference image. The viewpoint was shared between natural environments and gamut-rotated environments, but different camera angles were used for phase-scrambled environments (as the scene geometry was different). No tone-mapping was applied to any of the test images. Figure 5 shows 36 example test images together with an example lighting environment and its a*b* chromatic distribution. Mean luminance over the object region was 22.6 ± 3.44 cd/m^2^ (mean ± 1.0 S.E. across 36 objects). For each test image, the underlying physical reflectance parameters were defined as ground-truth values in the analysis.

**Figure 5:**
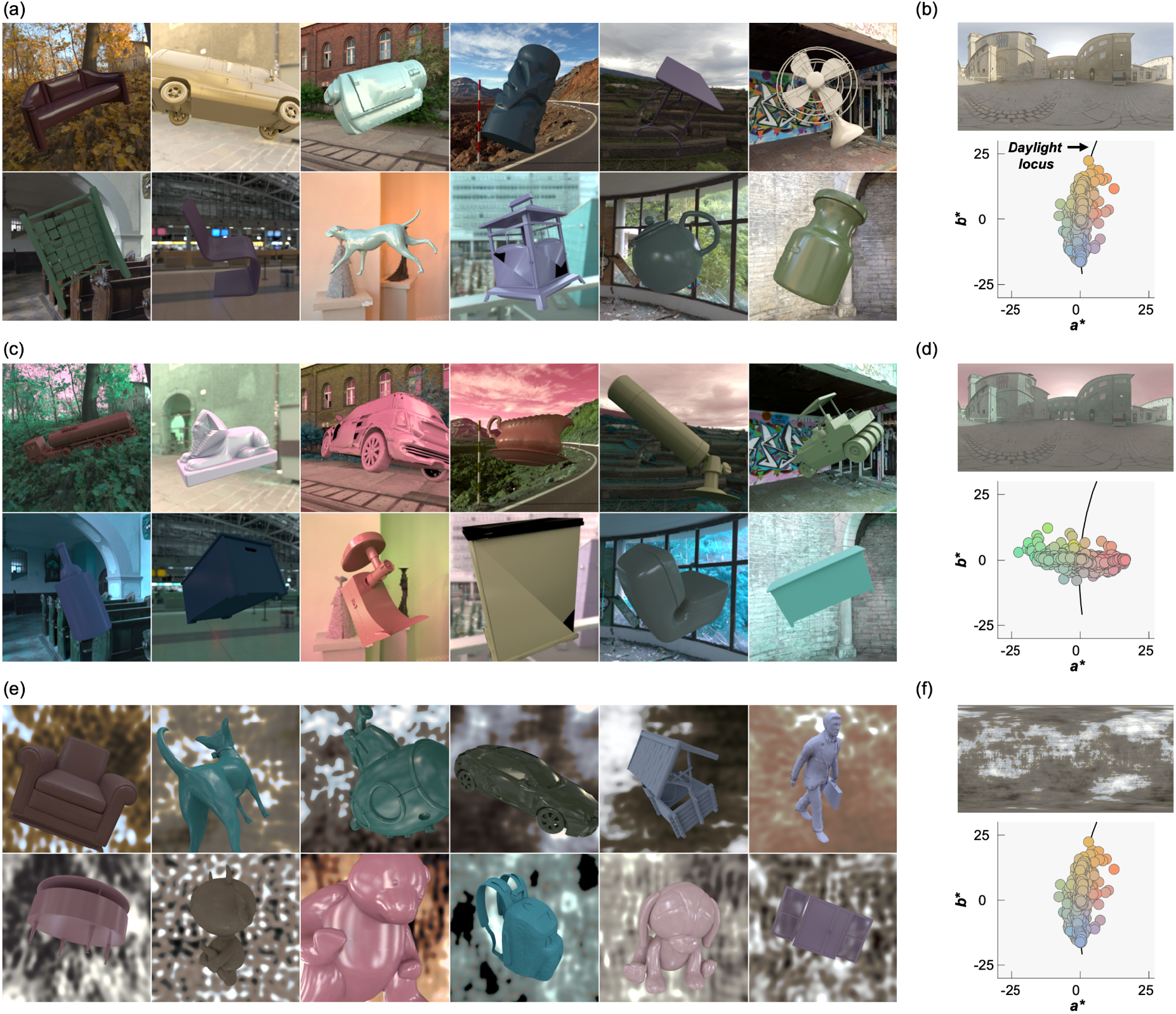
36 test images (a, c, and e) and an example lighting environment along with its *a*\**b*\* chromatic distribution (b, d, and f) in Experiment 1. (a, b) natural lighting environment, (c, d) +90° gamut-rotated lighting environment and (e, f) phase-scrambled lighting environment. Each test image contains a single object whose shape (and orientation), color and gloss level were randomly assigned.

For Experiment 2, we generated 216 test images, a factorial combination of 12 natural lighting environments and 18 shapes sampled from shapes used in Experiment 1. These images are shown in a later section for Experiment 2.

### 2.5 Procedure

Prior to the experiment, participants were given instructions both orally and by written text on the screen: *“Your task is to adjust glossiness and color of the right ‘reference’ object in terms of specularity, lightness, chroma, and hue until the reference object appears to be made of the same material as the left ‘target’ object. Each parameter changes the appearance of the reference object as shown here*.*”* Regarding the final sentence, we presented some example reference images similar to Figure 3 (c) to explain how changing each parameter influences the appearance of the reference object. Three practice trials immediately followed the instructions, using example test images that were not used in the actual experiment. During the practice trials, participants were asked to explore the full range of each parameter to familiarize themselves with the range of each parameter. After the practice trials, participants adapted for one minute to 20 Hz random-dot dynamic color noise whose mean chromaticity was equal to the chromaticity of equal energy white, then the experiment began. Initial physical reflectance parameters for the reference object were randomized for each trial. Viewing distance was kept constant at 49 cm from the LCD monitor. Participants signaled when the matching was completed by a button press and the selected underlying physical reflectance parameters of the reference object were recorded as the participant’s response.

In Experiment 1, there were 36 test images and 2 control images which contained the same bumpy sphere as the reference objects presented under the reference lighting environment (symmetric matching). The reflectance parameter in control images was fixed at hue 120, chroma 22, lightness 40, and *Pellacini’s* c 0.1305 for the control image 1, and hue 320, chroma 14, lightness 60, *Pellacini’s* c 0.0152 for the control image 2. One session thus consisted of 38 settings, and all participants completed 3 sessions in total. One setting took 66.4 ± 15.5, 53.5 ± 13.0, and 41.1 ± 7.9 seconds (mean ± 1.0 S.D. across 10 participants) for sessions 1, 2 and 3, respectively.

In Experiment 2, one session consisted of 216 trials (216 test images), and 2 sessions were completed. One setting lasted 11.5 ± 3.23 and 8.42 ± 2.18 seconds (mean ± 1.0 S.D. across 10 participants) for sessions 1 and 2, respectively.

For both experiments, there was a break between sessions.

## 3. Experiment 1

### 3.1 Results

#### 3.1.1 Control condition

We first show participants’ settings for the control image 1 (symmetric matching) in Figure 6. It is clear that participants can highly accurately set the reflectance parameters close to ground-truth values when the shape and lighting environment are identical between test and reference images. The accuracy of the setting was similar for the control image 2. This is not surprising but these error values can be usefully taken as a measure of matching precision for each parameter.

**Figure 6:**
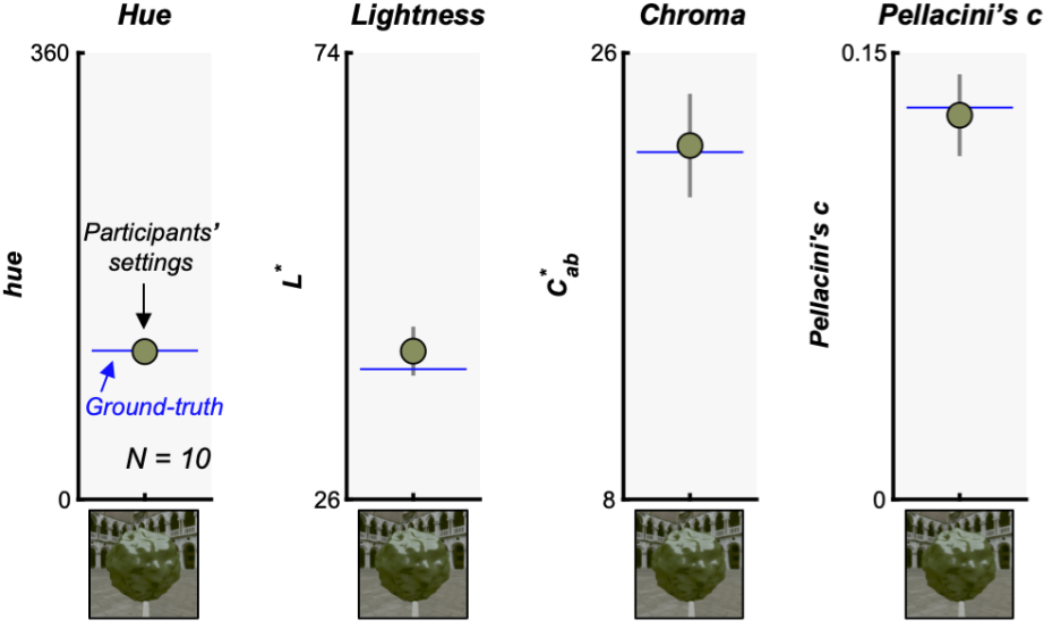
Results for the control condition (control image 1) where the test image and reference image had identical shapes and lighting environments to measure the precision of participants’ settings along each parameter. The settings were first averaged across 3 sessions for each observer and then averaged across 10 participants. The image on x-axis shows control image 1. The blue horizontal line depicts the ground-truth values assigned for the test object. The error bars show ± 1.0 S.D. across 10 participants.

#### 3.1.2 Main conditions

Figure 7 shows the main results for Experiment 1. Each data point shows the averaged setting across 10 participants for one test image. From left to right, each subplot shows results for hue, lightness, chroma and Pellacini’s c, respectively. Black numbers at the upper left corner of each subpanel show the correlation coefficient between ground-truth value and participants’ settings, calculated separately for each observer first and averaged across 10 participants.

**Figure 7:**
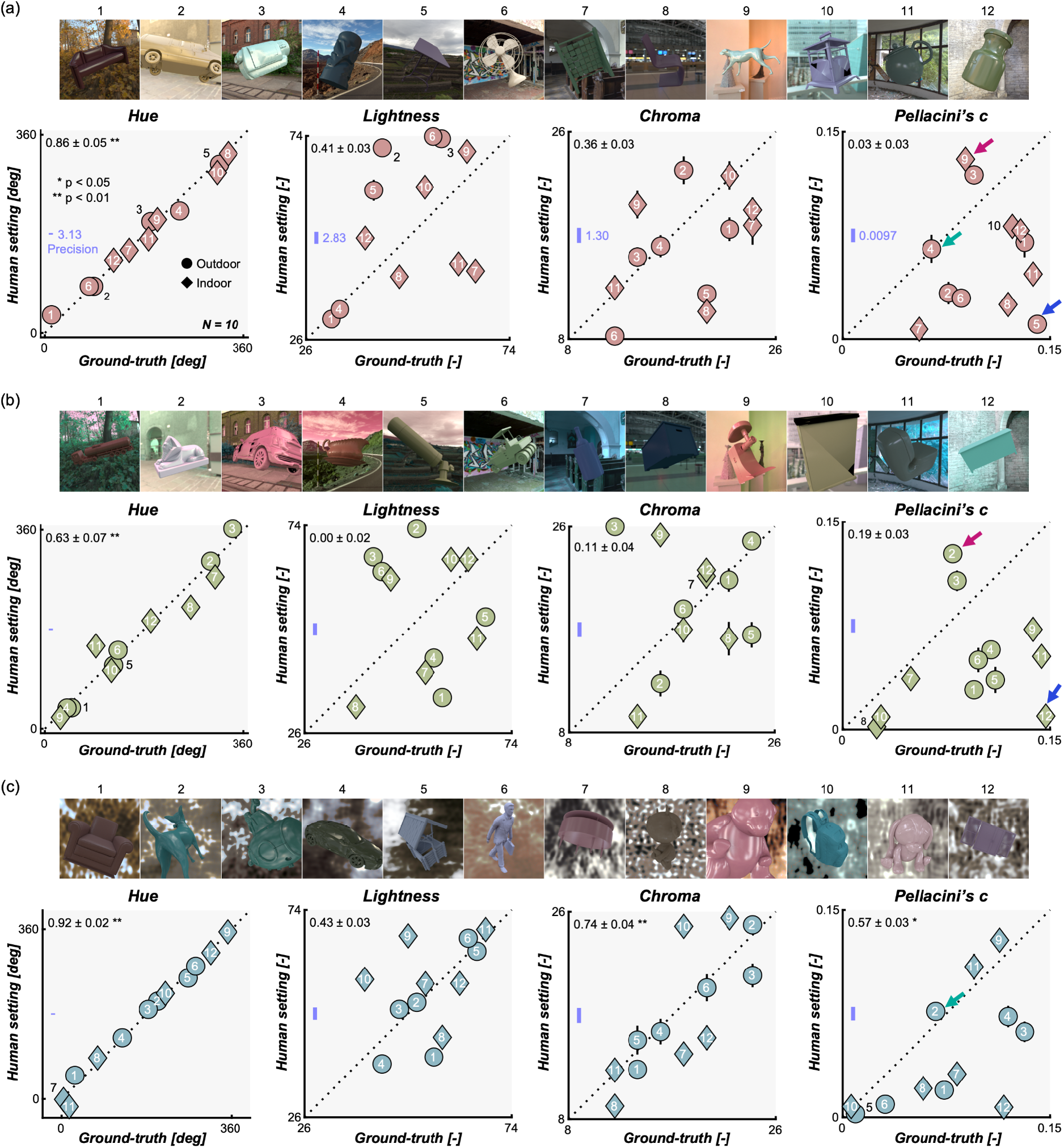
Results in Experiment 1 for (a) natural lighting environments, (b) gamut-rotated lighting environments, and (c) phase-scrambled lighting environments. Each setting was averaged across 10 participants. Error bars show ± 1.0 S.E.M. over 10 participants, which is smaller than the data point for most cases. Upper images in each panel show test images, with numbers showing the correspondence to data points. The number at the left-upper corner in each sub-panel shows the correlation coefficient (mean ± 1.0 S.E.) between ground-truth values and participants’ settings calculated separately for each participant and averaged across 10 participants. The blue line represents the precision computed from the mean absolute error in the symmetric matching data. Small colored arrows show test images whose gloss level was judged to be particularly high (red arrows), low (blue arrows) and the same level as the reference image (green arrows).

Note that in an asymmetric matching experiment participants’ settings should be interpreted relative to the reference image used in this study. Taking the Pellacini’s c as an example, data points falling exactly on the diagonal unity line shows that participants set the Pellacini’s c of the reference object so that it is the same as the test object. This means that the perceived gloss level of the test object and reference objects are the same when both objects have the same Pellacini’s c. In an alternative case, data points located below the diagonal line mean that those test objects would appear less glossy than the reference object if the test objects’ Pellacini’s c (ground-truth) were applied to the reference object (which is why participants needed to lower the Pellacini’s c of the reference object to achieve matching). For this reason, absolute error values are likely to vary by the choice of reference image, and thus we evaluated the accuracy of the settings using correlation coefficients between human settings and ground-truth values which are not affected by additive shift or multiplicative scaling of the data points which might have been introduced by the choice of the reference image. This relative measure also allows comparison across different parameters.

At first glance, data points in Figure 7 show large scatter, except for hue settings, and the data seem noisy. However, we emphasize that these settings were highly consistent across participants as illustrated in Figure 8, meaning that all participants systematically showed similar deviations from ground truth. Left bars show the averaged correlation coefficient between human settings and ground-truth (the same as the black numbers in the upper left corners of plots in Figure 7), and small circles show individual participants. Right bars show the averaged correlation among participants (inter-participant correlation). To compute this, for each participant, we calculated the correlation coefficient between the participant’s settings and settings averaged across all participants (Nili et al., 2014). This reveals that human-human correlations are generally high regardless of human-groundtruth correlation, making it clear that the scattered data patterns in Figure 7 are not due to noise.

**Figure 8:**
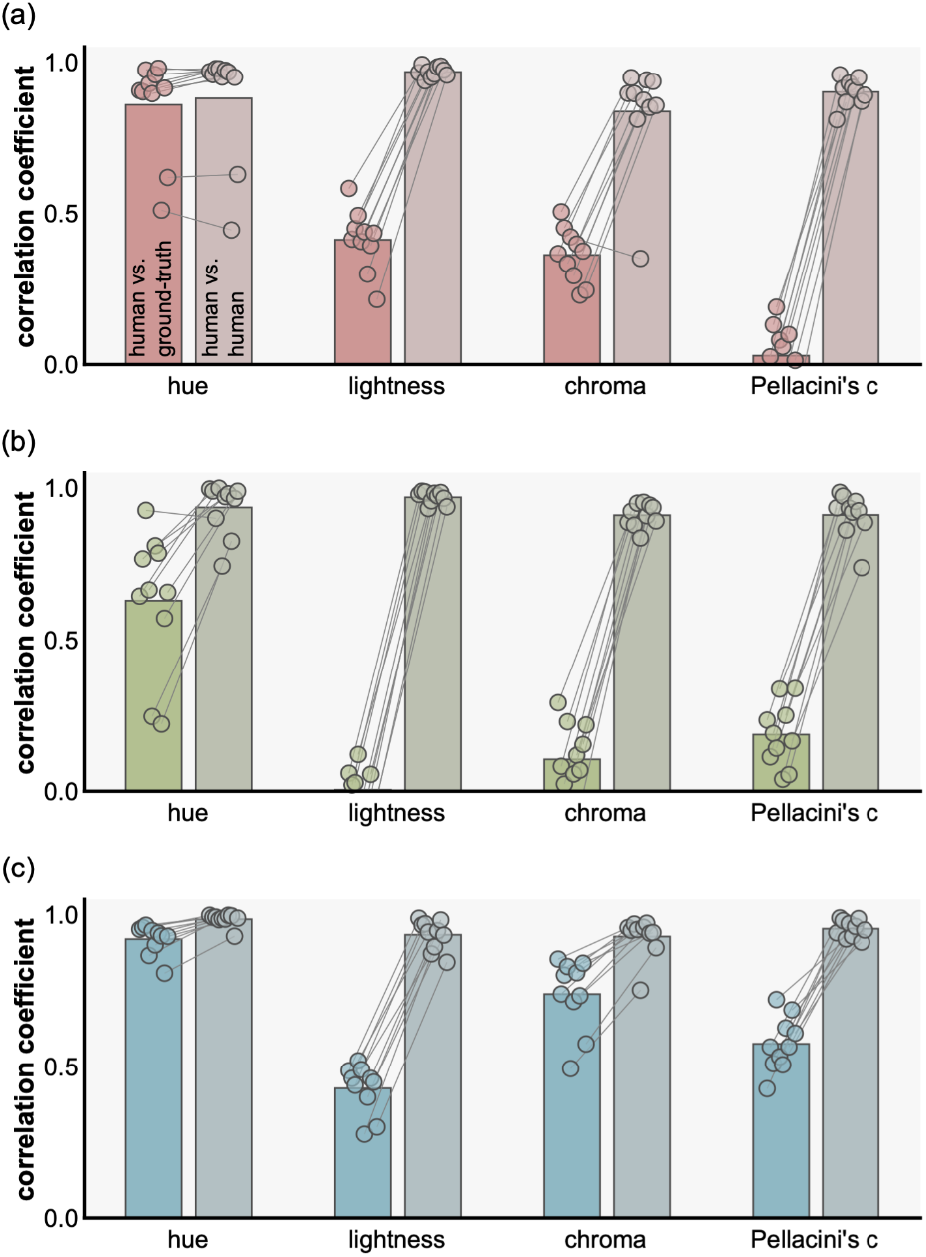
Left bars and data points show correlation coefficients between each human participant and the ground-truth and right bars and data points show inter-participant correlation (correlation between each participant and the average across all participants) for (a) natural, (b) gamut-rotated, and (c) phase-scrambled lighting environments. Each dot shows an individual participant and each bar shows the average across 10 participants.

Returning to Figure 7, hue settings in (a) natural environments show a significant correlation with ground-truth values, indicating that hue judgment is stable regardless of test lighting environments. This observation holds well for (c) phase-scrambled environments and less so for (b) gamut-rotated environments. This generally high degree of hue constancy may not be very surprising because the circular correlation between mean hue value over the object region and ground-truth values across 36 test images was 0.92. This means that basing a judgment on the mean hue value over the object region (Milojevic et al. 2018) in each trial would lead to high correlation with ground-truth, but this might also suggest that hue is physically a relatively stable quantity at least for the environmental illuminations we tested here. To evaluate the influence of the type of lighting environment (natural, gamut-rotated, and phase-scrambled) on the correlation coefficient between human settings and ground-truth, we performed one-way repeated measures ANOVA which confirmed a significant effect of illuminant type (*F*(2,18) = 11.9, *p* = 5.14 × 10^−4^). Post-hoc multiple comparisons (Bonferroni’s corrected *p* = 0.05) showed significantly higher correlation for (a) natural environments than (b) gamut-rotated environments and (c) phase-scrambled than (b) gamut-rotated environments, suggesting worse hue constancy in chromatically atypical lighting environments which implies the role of a daylight prior in judging the illuminant influence (Pearce et al. 2014; Weiss, Witzel & Gegenfurtner 2017).

In contrast, lightness and chroma settings are scattered in a disorderly way, leading to generally lower correlation with ground-truth than for hue settings. The correlation coefficient was statistically significant only for the chroma setting in (c) phase-scrambled lighting environments. Similarly, settings of Pellacini’s c were not highly correlated with ground-truth though the correlation is significant for (c) phase-scrambled environments. Here most data points fall below the diagonal unit line, meaning that the perceived gloss level of test objects was generally lower than the reference object. This might reflect either the more even sampling of surface normals for the reference geometry, or a tendency of the Uffizi probe to make objects appear particularly glossy.

### 3.2 Discussion

Subsequent subsections discuss potential underlying reasons for highly consistent error patterns for chroma, lightness and gloss judgments.

#### 3.2.1 Interaction between diffuse reflectance and specular reflectance

Firstly, we considered the possibility that specular reflections may have contaminated the perception of body color. If so, we should observe that deviations between lightness and chroma settings by participants and ground-truth values become larger as a function of physical (or perceived) gloss level. Accordingly, we computed correlation coefficients between the ground-truth Pellacini’s c and lightness error (human setting minus ground-truth) and between the ground-truth Pellacini’s c and chroma error across 12 test images, separately for each type of lighting environment. However, we found no significant correlation under any type of lighting environment. We repeated this analysis using Pellacini’s c set by participants (i.e. perceived gloss) instead of the ground-truth Pellacini’s c, but again there was no significant correlation under any type of lighting environment. Thus, large errors in chroma and lightness observed in Figure 7 are unlikely to be due to masking or intrusion by the specular reflection.

Similarly, we next asked whether diffuse reflection contaminated the perception of gloss. In other words, were participants more likely to make errors in gloss settings for a certain range of body colors? Figure 9 visualizes the magnitude of errors (Pellacini’s c set by participants minus ground-truth Pellacini’s c) as a function of body color. However, we found no significant correlation between errors in gloss settings and each of the color parameters (i.e. hue, chroma and lightness) under any type of lighting environment after the correction of significance level (by Bonferroni’s correction), showing that there are no noteworthy interactions between failures of gloss constancy and body color.

**Figure 9:**
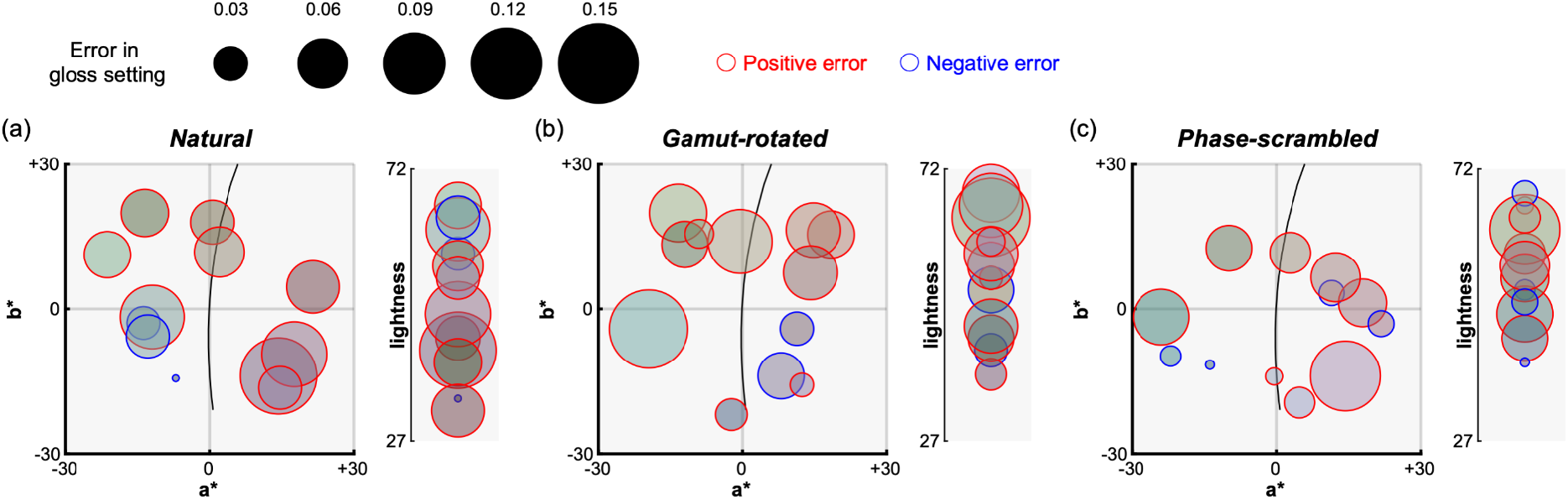
Analysis of interaction between body color (diffuse reflectance) and perceived gloss level for (a) natural, (b) gamut-rotated, and (c) phase-scrambled lighting environments in Experiment 1. Each data point corresponds to one test image and its color shows the body color (in sRGB format). The size of each data point represents the error of gloss setting (human setting minus ground-truth). Red and blue edge colors show positive and negative errors, respectively. Left plot shows representation in *a*\**b*\* chromatic plane while the right plot shows lightness axis. There was no systematic trend found between the body color and the magnitude of gloss errors.

Overall, these analyses confirmed that error patterns in color settings were independent from specular reflectance. Similarly errors in gloss judgements were not systematically affected by diffuse reflectance. Evidently judgments of the components of reflectance are based on distinct image information or internal representations, not a single composite representation of the entire bidirectional reflectance distribution function (BRDF; Nicodemus et al. 1977).

#### 3.2.2 Interaction between lightness and chroma

We wondered whether there might be an interaction between parameters related to color, which affected participants’ perceptual judgment and matching. One such candidate was the interaction between chroma and lightness as a past observation suggested that perceived saturation is relatively well predicted by *C*_*ab*_* / *L*\* (Fairchild, 2013; Schiller et al. 2018). In other words, it is possible that participants judged the chroma and lightness match by simply judging the match in perceived saturation between test and reference objects. If that’s the case, ground-truth values and participants’ settings when represented in *C*_*ab*_* / *L*\* should correlate well. As shown in Figure 10, we found significant correlations for all three types of lighting environment. This explains at least partially the scattered setting patterns for chroma and lightness in Figure 7, especially for the (c) phase-scrambled environments, although a substantial portion of the variance remains unaccounted for with the (a) natural and (b) gamut-rotated environments

**Figure 10:**
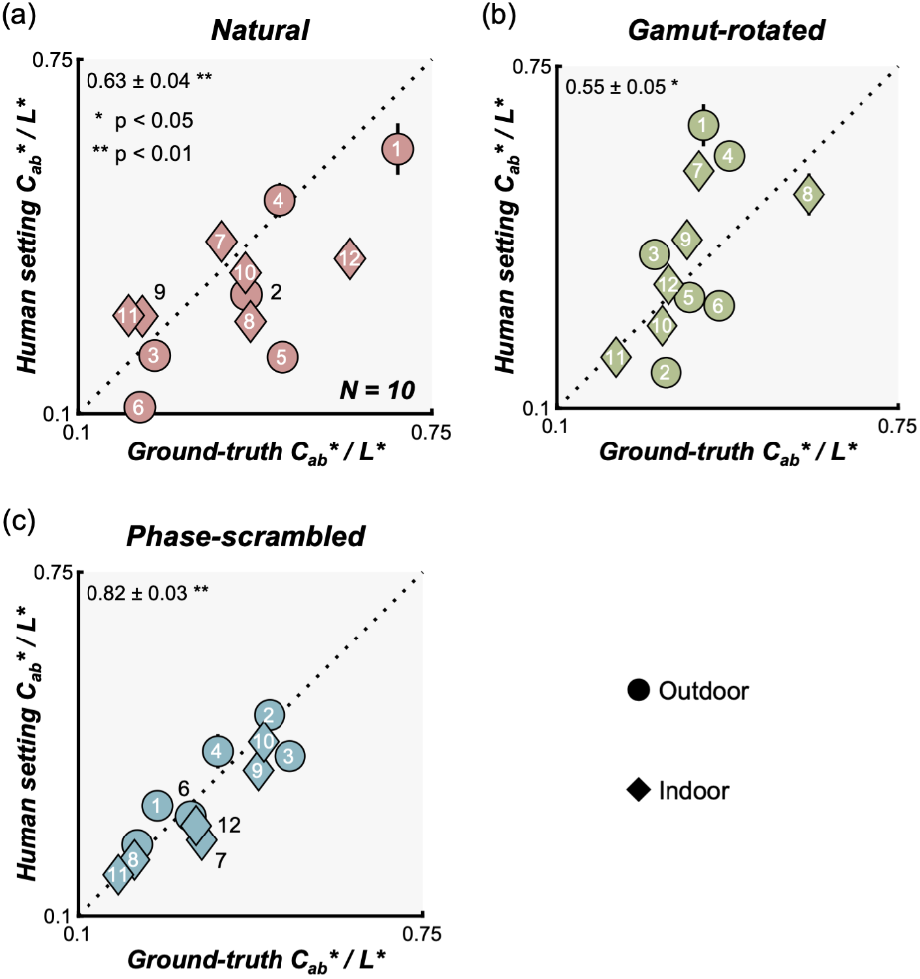
Scatter plot to see whether interaction between chroma and lightness (saturation defined as *C*\*_*ab*_ / *L*\*) can explain the observed chroma and lightness constancy failures. The x-axis shows ground-truth saturation while the y-axis shows saturation computed from matching results. Significant correlations between ground-truth and human settings suggest that participants might have used the perceived saturation to help determine whether the color was matched between reference object and test object.

#### 3.2.3 Image statistics

Another candidate account would be that participants based their judgements on various summary statistics directly accessible from test images. Indeed, since there is no direct way for participants to access the ground-truth values of gloss and body color, such a strategy could be a reasonable alternative approach to performing the task. To predict participants’ lightness and chroma settings, we calculated mean, median, standard deviation, skewness, kurtosis, first quartile (Q1), third quantile (Q3), minimum and maximum values of lightness and chroma across the object region in each test image. Surrounding context was excluded from the computation.

Additionally, to understand why gloss constancy was poor in our task, we selectively looked at test images that were rated particularly high (discrepancy from ground-truth, +0.0411, +0.0469) and low gloss (discrepancy, -0.129, -0.136) as well as images where ground-truth and human settings well matched. Figure 11 (a) shows example images (these images are labeled by colored arrows in Figure 7). It is evident that specular reflection patterns are visibly different across images. Objects in high gloss images (surrounded by red squares) seem to receive strong directional lights in the environment and consequently have a readily visible specular reflection pattern though physically the specularity is around the middle of the range (see Figure 7). In contrast, objects in low gloss images (surrounded by blue squares) are placed under a dim and cloudy lighting environment or indoor scene. Specular reflection is present in these cases too, but it spreads across the surface and that’s why when mixed with diffuse images, the specular reflection patterns are hard to detect even though both objects have nearly the highest possible specular reflectance. Finally, for the images where human settings and ground-truth were well correlated, specular reflections are moderately visible. In sum it visually makes sense that these images were rated by participants in these ways, which encouraged us to compute image metrics based on patterns of specular reflection.

**Figure 11:**
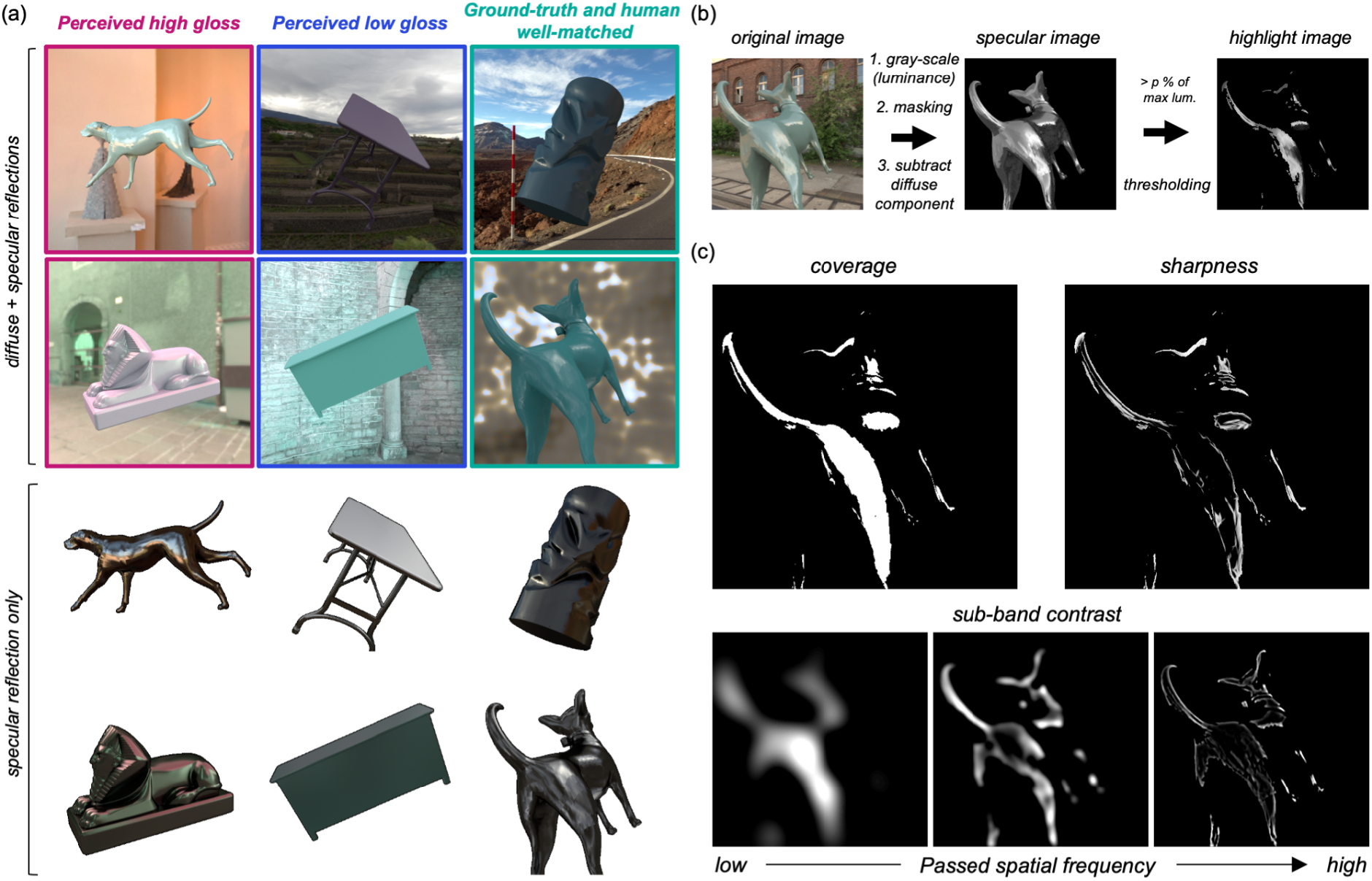
(a) Example images where humans perceived high gloss (left column), low gloss (center column) and where ground-truth gloss level and human settings matched well (right column). Upper 6 images show original images where diffuse and specular reflections are both included. Lower 6 images show objects without diffuse reflection and surrounding context to visualize the spatial structure of specular reflection on which participants might have based their gloss judgements. (b) Process to convert original test image to thresholded highlight image. (c) Calculation of three metrics to predict human gloss percepts: coverage, sharpness and sub-band contrast (see main text for details).

Thus, in addition to basic descriptive statistics, we used metrics computed from the structure of the specular reflection image (Marlow et al. 2012; Schmid et al. 2021) to predict participants’ gloss settings. As shown in Figure 11 (b), we first converted the original image to a luminance image, masked out the object region, and subtracted the diffuse component from the image, which resulted in test images with specular reflection alone. Then, we extracted pixels whose intensity is higher than *k* % value of the highest intensity across this specular image, where *k* took the following values 0, 1, 3, 5, 10, 20 and 40 to get rid of the region of specular reflection that stems from secondary and higher-order inter-reflections. Using this thresholded ‘highlight image’ we calculated the following three metrics. The first metric is *coverage*, corresponding to the proportion of area covered by the highlight relative to the whole object area as depicted in the upper-left part of panel (c). Second, we calculated the *sharpness*. Using a spatial convolution, this metric emphasizes the region where luminance rapidly changes and sharpness is defined as a mean value of the convoluted sharpness map (Vu et al. 2012) as shown in upper-right part of panel (c). For coverage and sharpness, model predictions were affected by the cut-off percentage to threshold the highlight regions and thus we selected an optimal value of *k* that produced the highest correlation value with human settings. Finally, the third metric was *contrast*, which essentially measures the spatial luminance variation over the surface. The standard way would be to calculate a contrast from the raw highlight image directly. However, considering a previous observation that perceived gloss is affected by the modulation of a specific frequency channel (Boyadzhiev et al. 2015), we first decomposed the raw highlight image into 8 sub-band images using a Gaussian band-pass filter (upper and lower cut-off frequencies: 1.5 to 3.0, 3.0 to 6.0, 6.0 to 12.0, 12.0 to 24.0, 24.0 to 48.0, 48.0 to 96.0, 96.0 to 192, and 192 to 384 cycles/image) and a subset of sub-band images are shown in the lower part of panel (c). We calculated the RMS contrast, equivalent to standard deviation of the pixel intensities, for each sub-band image as well as for an aggregated image across all frequencies. This means that unlike coverage and sharpness which has one parameter, the sub-band contrast metric has two free parameters (i.e. cut-off pixel intensity and cut-off spatial frequency band), and optimal values producing the highest correlation with human settings were selected. For all three metrics, searching for the best parameters was performed separately for each type of lighting environment (natural, gamut-rotated, phase-scrambled).

Each bar in Figure 12 shows the correlation coefficient between image statistics and human settings for (a) natural lighting environment, (b) gamut-rotated lighting environment, and (c) phase-scrambled lighting environment. Higher values indicate that the models capture participants’ settings better. The magenta shaded region shows the inter-participant noise ceiling, between upper bounds (same as magenta values in Figure 7) and lower bounds, computed by a similar procedure as the upper bound, but by calculating correlation coefficient between the left-out participant’s settings and settings averaged across all other participants. This range effectively defines a bound on how well any image-computable model could perform. These image statistics were computed from test objects that include both diffuse and specular reflections. For lightness and chroma, we also considered image statistics computed directly from the diffuse component only (without specular reflection) to test the idea that humans might have effectively discounted the specular reflection from the test image. The red diamonds show the correlation between image statistics calculated directly from the diffuse image and participants’ settings. If participants took such a strategy, the red diamonds should come higher than the green bars.

**Figure 12:**
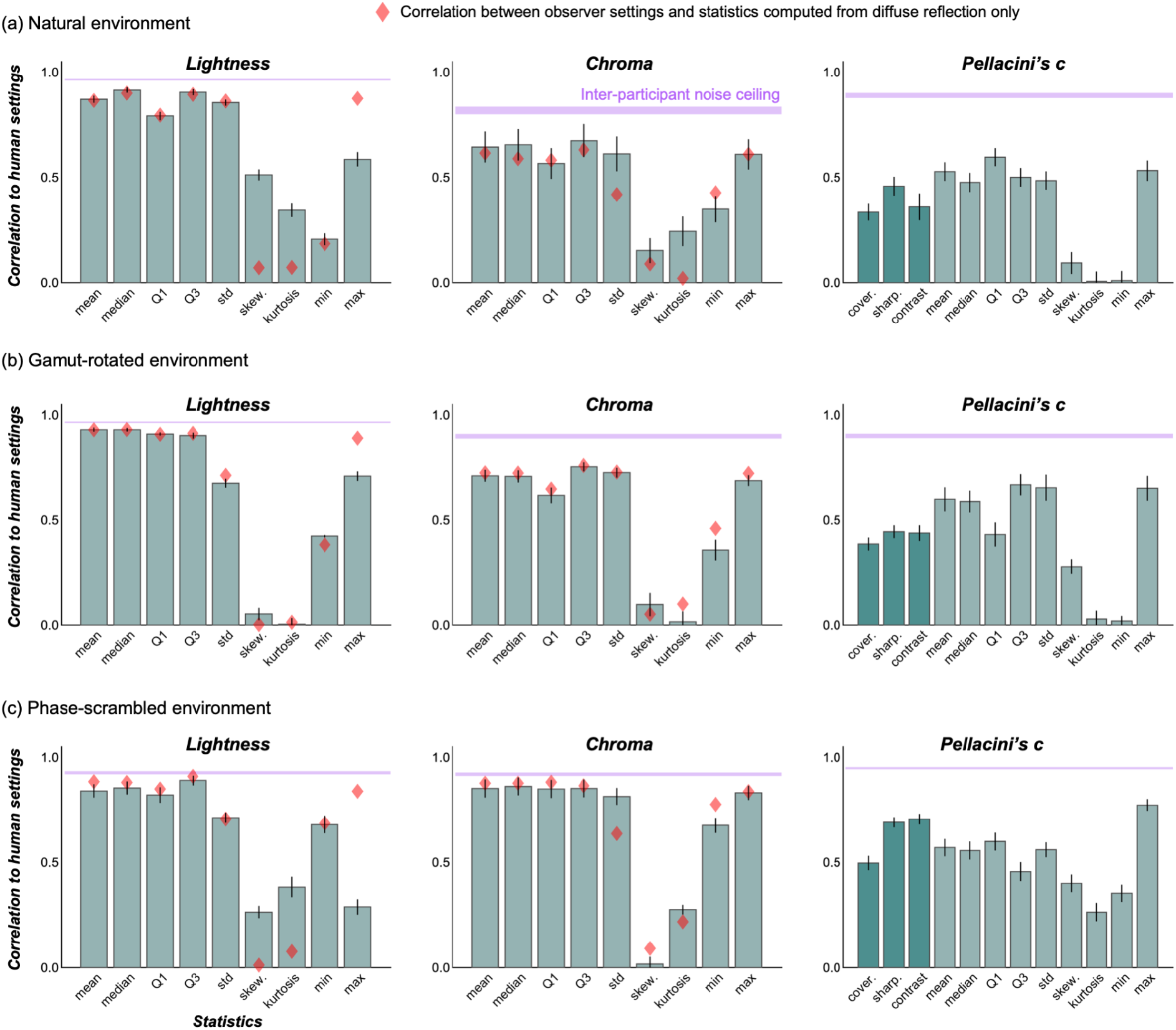
Correlation coefficients between image statistics (calculated over the object regions in test images) and human settings. (a) Natural environment, (b) gamut-rotated environment and (c) phase-scrambled environment. The magenta shaded areas show an inter-participant noise ceiling, equivalent to the correlation across participants, which correspond to magenta numbers in Figure 8. Red diamonds show the correlation between image statistics calculated over the object region in test images that had only diffuse reflection (no specular reflection).

For lightness and chroma, it is clear that although the correlation between human settings and ground truth are remarkably low (as shown in Figure 7), participant’s settings are highly correlated with simple statistics such as mean lightness and mean chroma over the object region, nearly touching the noise ceiling level in some cases. Interestingly, for lightness, maximum luminance of diffuse components (red diamonds) predicts human settings better than maximum luminance of the original image (bars), consistent with a past observation (Giesel & Gegenfurtner, 2010).

Looking at Pellacini’s c, overall image statistics correlate with human settings less than for chroma and lightness, and maximum luminance values are generally good predictors for any type of lighting environment. Yet, there are still significant disparities between the predictors and human performance. Thus these models explain the failure of gloss constancy to some extent, but not enough to capture the complexity of gloss perception.

To summarize the results from Experiment 1, both success and failures of color constancy were well captured. Hue constancy holds remarkably well. Failures of chroma and lightness constancy were systematic and largely predicted by simple metrics such as mean chroma or mean lightness over the object surface. Gloss constancy was very poor for our stimuli, but matches were highly consistent across participants, and the simple image statistics we investigated explained human behavior to a limited extent.

To grasp the complex nature of gloss perception better, we felt that the 36 test images used in Experiment 1 may not be enough. Thus we conducted a follow-up Experiment 2 with 216 test images using a factorial combination between 12 lighting environments and 18 shapes, and the perceived gloss was again measured using the asymmetric matching task.

## 4. Experiment 2

### 4.1 Test images and procedure

We generated 216 images using 18 shapes sampled from those used in Experiment 1, and 12 lighting environments (Figure 13). Hue, lightness and chroma of test objects and matching objects were fixed at 188.2°, 33.9 and 11.9, respectively. We picked this greenish color as we speculated that visibility of specular reflection would be higher when the body color falls on the axis orthogonal to daylight locus (i.e. red-green axis) because as seen in Figure 4 colors of specular reflection are mainly distributed along the blue-yellow axis. The task of participants and procedure were identical to Experiment 1, except that participants’ adjusted only Pellacini’s c in Experiment 2 and the color of the reference object was fixed to the same green as the test object. For each test image, we applied a random Pellacini’s c ranging from 0 to 0.224, which corresponds to the range from 0 to 0.0999 in Ward’s specularity. The maximum value of Pellacini’s c here differs from the value in Experiment 1 (0.149) because the lightness value of the object for the conversion between Pellacini’s c and Ward’s specularity was 33.9 instead of 50. One session consisted of 216 trials, and all participants completed two sessions in total.

**Figure 13:**
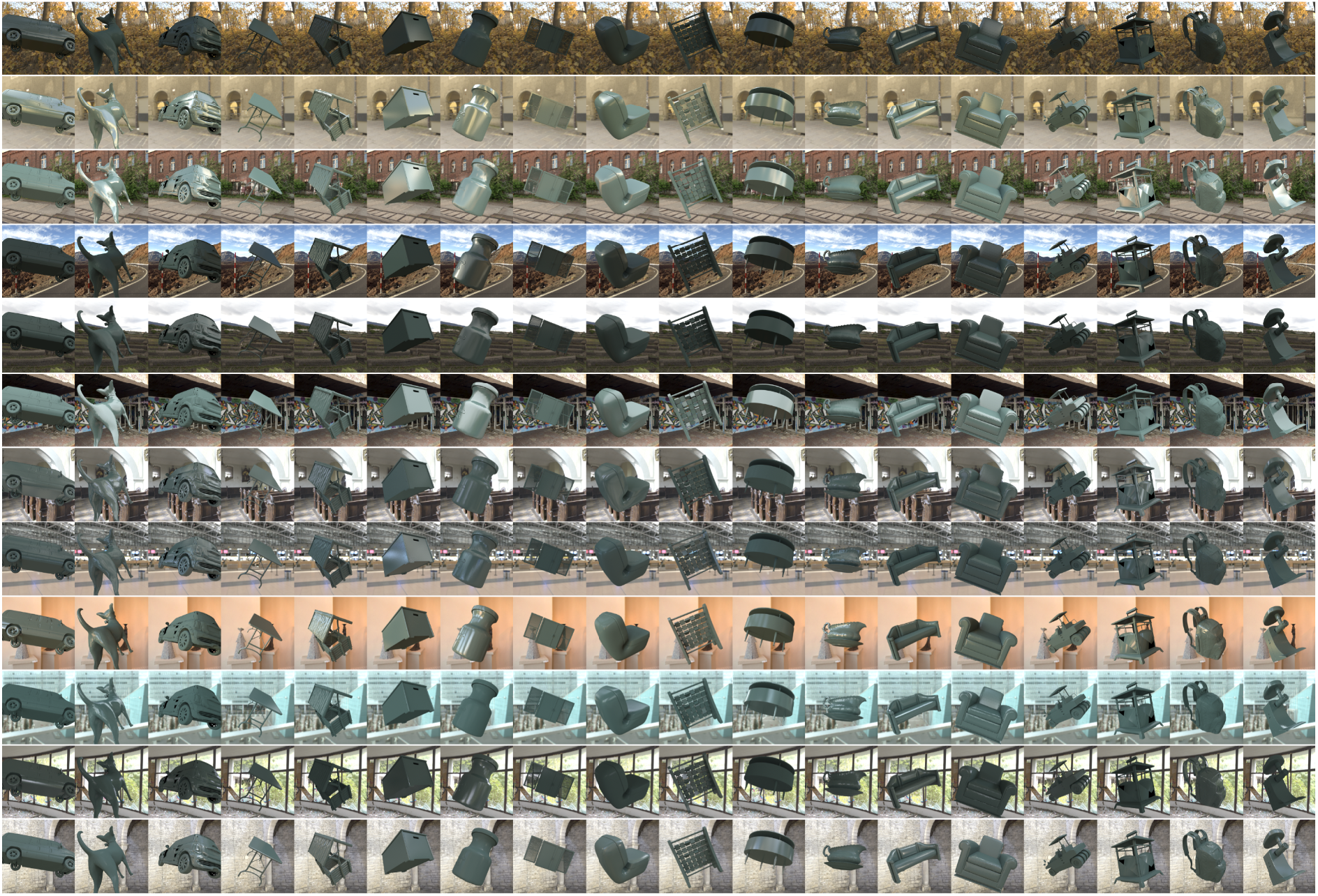
216 test images used in Experiment 2. Images were generated by the factorial combination between 12 natural lighting environments in Experiment 1 and 18 shapes that were randomly sampled from the 36 shapes in Experiment 1. Body color was fixed as a greenish color, which we found useful to increase the visibility of specular reflection on the objects’ surface.

### 4.2 Results and Discussion

Figure 14 (a) shows participants’ settings (averaged across 10 participants) in Experiment 2 grouped by the test lighting environment. Each data point corresponds to one shape, and thus there are 18 data points in each subplot. Globally looking through subplots, it is evident that data points deviate substantially from the unity line (diagonal dotted line), showing that human settings and ground-truth strongly disagree. It is also noticeable that the slope of the red regression line differs from one lighting environment to another (min 0.34 and max 0.90). Higher slope means that on average under that lighting environment objects appear more glossy. Also, the correlation coefficients between human settings and ground-truth values (black numbers at upper-left) were significant in all lighting environments, unlike Experiment 1, but the values vary from 0.46 to 0.78 showing that the variability due to the objects’ shape also differs from one lighting environment to another. Thus, simple transformations are unlikely to equate perceived gloss level in one lighting environment to another environment. This is inconsistent with past research using smooth spheres (Fleming et al, 2003; Doerschner et al. 2010), and the use of a variety of shapes in this study is likely to be a reason. Overall, these results suggest that perceived gloss somewhat correlates with underlying physical specular reflectance, but also differs due to the change of lighting environment, showing failures of gloss constancy.

**Figure 14:**
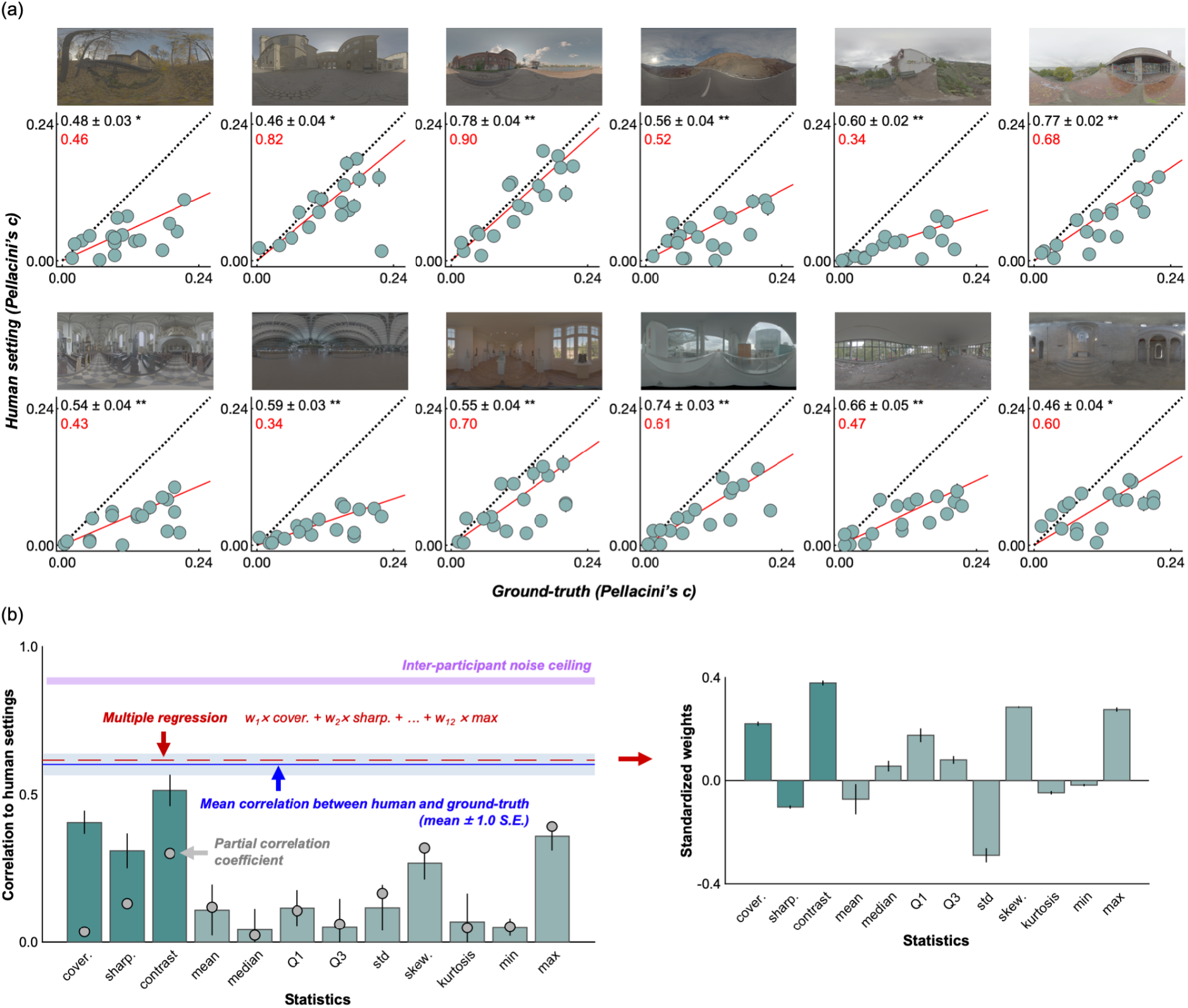
(a) Average setting across 10 participants in Experiment 2, where settings were grouped by lighting environment. Each data point denotes one shape, and thus there are 18 data points in each scatter plot. Error bars show ± 1.0 S.E. across 10 participants. Regression lines fitted to the 18 data points are shown by a solid red line. Black numbers in the upper left corners of the plots show the correlation coefficient (± 1.0 S.E.) between human settings and ground-truth, computed for each participant first and then averaged across 10 participants. Red numbers show the slope of the regression line. Higher slope means that, on average, that lighting environment produced a high degree of perceived gloss. (b) Lefthand bar plot shows correlation coefficient between image statistics and human settings. The dark green bars show image statistics computed from specular reflection images, and the light green bars show simple luminance image statistics. Error bars show ± 1.0 S.D across 10 participants. The blue line shows the correlation coefficient between participants’ settings and ground-truth values, averaged over 12 lighting environments (average over 12 black numbers in panel (a)). The pink shaded areas show an inter-participant noise ceiling computed in the same way as for Experiment 1 (Figure 12), which no model can exceed. The gray filled circles show the partial correlations between each image statistic and human settings, removing the influence of the correlation between image statistics and ground-truth. The red line shows the correlation coefficient between human settings and a weighted sum of 12 statistics where weights were optimized through multiple regression. Right-hand bar plot shows a standardized regression coefficient (Borgonovo & Plischke, 2016) for each statistic averaged across 10 cross-validations, and error bars show ± 1.0 S.D.

We next asked whether image statistics explain these human behaviors. The lefthand bar graph in Figure 14 (b) shows the correlation coefficient (averaged over 10 participants) between each statistic and human settings over 216 images. Unlike the results of Experiment 1 (Figure 12), basic image statistics showed fairly low correlation, while the sub-band contrast metric showed relatively high correlation. However, this correlation was significantly lower than the correlation between human settings and ground-truth shown by the blue line (two-tailed paired t-test; *t*(18)=3.25, *p* = 0.0045). Moreover, it is worth noting that this contrast metric (and others) also correlates with ground-truth value to some extent, and this may be a part of the reason why these metrics correlated with human settings. Thus, to remove the influence of ground-truth, we additionally computed a partial correlation coefficient between the image statistics and human settings (gray filled circles), removing the correlation between image statistics and ground-truth values. We found that they are lower than the original correlation coefficient especially for image metrics computed from specular reflections (left three gray points). This result suggests that these models correlate well with human settings mainly simply because they correlate with ground-truth, not necessarily capturing systematic failures of gloss constancy consistently exhibited by participants. For other simple image statistics, partial correlation and original correlation coefficients are similar to each other, likely because those simple metrics do not correlate with ground-truth values well. Finally, we tested what happens if we assume observers combined multiple statistics instead of using a single cue independently. To test this idea we ran multiple regression analysis using all 12 metrics as independent variables to explain the participants’ settings. When we did this, we trained a regressor using the averaged settings across 9 participants to find optimized weights for 12 metrics, and then using these optimized weights we calculated the correlation between the regressor’s prediction and the settings by the left-out participant. We repeated this 10 times (leave-one-out cross-validation), and the average across the 10 correlation coefficients is shown by the horizontal dashed red line. The right-hand bar plot shows standardized weights for each statistic, averaged across 10 validations. However, the improvement due to integration of multiple cues was marginal, and consequently there is still much room between this level and the noise ceiling level.

## 5. General Discussion

How do we overcome huge variations in the proximal image to create a stable percept of the color and gloss of objects? To address this question we measured color and gloss constancy together using an asymmetric matching task under a diverse set of lighting environments. Our results revealed a strong asymmetry across hue, chroma and lightness constancy; the degree of hue constancy was generally high, although it slightly decreased when the lighting environment had atypical chromatic properties, while lightness and chroma constancy were in general severely limited (except for chroma settings under phase-scrambled environments). These failures of chroma and lightness constancy were well captured by the saturation metric (chroma/lightness) and simple image statistics over the object region in the image. In contrast, gloss constancy was generally poor (i.e., gloss ratings depended on lighting environment and shape), but phase-scrambling directional lighting geometry did not additionally impair gloss constancy. Image statistics explained those failures of gloss constancy only to a limited extent. One major finding in this study is that, although there have been observations that simplistic image metrics can account for large variations in human gloss perception, when the diversity of shape and lighting approaches that seen in real world environments a significant amount of consistent variance in gloss judgments remains unexplained.

This study presented constancy errors that were remarkably consistent across tested participants, but there have been reports that the degree of color constancy could vary substantially from one experimental condition and paradigm to another (Foster 2011). Our findings are in fact largely in line with findings from several past studies. Pont & te Pas (2006) reported that participants showed significant failures of constancy in a task where two presented spheres illuminated differently have either the same reflectance properties or not. Nishida & Shinya (1998) used a reflectance matching paradigm using both albedo and specular reflectance and showed that participants’ lightness and glossiness judgment were heavily influenced by object shape. Olkkonen and Brainard (2010) showed systematic failures of gloss constancy due to changes of lighting environment. However, it is worth noting that all these experiments reporting constancy failures—including our own—were conducted using computer monitors, and whether this finding applies to real-world scenarios needs careful investigation. For example, one challenge associated with performing the asymmetric matching paradigm on monitors is limited adaptation to test lighting environments, which is a key contributor in human color constancy (Smithson, 2004). Although the participants were allowed to move their eyes freely during a trial, it is reasonable to assume that the participant looked at the reference image for most of the time during the trial to complete the task. Moreover, images presented on the monitor only occupied a small part of the visual field, and the surrounding region in test images that provides cues to the lighting environments was even smaller (on average, 26.3 % of the whole image for Experiment 1 and 18.3 % for Experiment 2), which would have made it harder for observers to infer the illuminant influence. However, it is worth noting that the conditions in our experiment were sufficient to enable relatively good hue matches. Finally, we presented only one object in each test image, and presenting multiple objects with various colors in the same scene may have increased the degree of color constancy.

We observed that changing the lighting environment had different effects on different color dimensions. One reason for the superior hue stability could be that the lighting environments we selected did not produce extreme chromatic shifts in the proximal image and consequently pixel hue values did not change severely enough to cause poor constancy. In fact, as shown in Figure 4, although we tried to select environmental illuminations whose color distributions are different from each other, the mean color (shown by a black cross symbol) is still located relatively close to the white point. In contrast, chroma and lightness values shifted enough that mean chroma and lightness values were decorrelated from ground-truth values. However, it is generally true that in natural environments extreme chromatic shifts are rare (Morimoto et al. 2022). It is thus interesting to ask how much our selection of lighting environments reflects the true variation of actual lighting environments. If the physical hue values of an object do not vary substantially across scenes in the real world, hue serves as a useful perceptual anchor for object identification under different environments (Milojevic et al. 2018; Ennis 2018).

We showed that a contrast metric computed directly from the specular images showed highest correlation with human settings in Experiment 2, but this model implicitly assumes that humans are capable of separating diffuse and specular reflections from a given image. Such a separation is an ill-posed problem and it is an empirical question how accurately humans can perform the decomposition (e.g. Lee & Smithson 2018). A complete model of human gloss perception should predict the perceived gloss level from a raw image where diffuse and specular reflections are confounded. A recent effort used deep neural networks trained to output a specular image from a raw image and showed that such a network outperformed a simple alternative highlight-detection model based on thresholding and showed relatively high overall similarity to human judgements (Prokott et al. 2022). Another trained unsupervised deep neural networks to model the high-level statistics of images of glossy and matte surfaces, and found that these predicted human gloss judgements better than supervised networks or a range of simpler image statistics (Storrs et al., 2021).

A good perceptual model should reproduce both the successes and error patterns that humans make on an image-by-image basis beyond predicting the overall performance level (Geirhos et al. 2020; Storrs et al., 2021). In this sense, systematic error patterns in Experiment 2 are a potentially useful feature of the dataset as a window into underlying constancy mechanisms. However, we found that our hand-selected features accounted for a limited extent for gloss percepts, and it is a common shortcoming that researchers must select in advance or hand-engineer candidate features. In recent years ‘big data’ approaches (often coupled with deep neural networks) have been opening a new avenue to overcome such limitations as networks can learn to extract useful image features by themselves (Prokott et al. 2021; Tamura et al. 2021; Liao, 2022; Sawayama, 2022), and this study might also benefit from such approaches. The fact that human judgments can deviate substantially but consistently from ground truth—as we found here—suggests that training a neural network with human responses would potentially yield quite different internal representations than training with ground truth specular reflectance values.

## Acknowledgement

Authors thank Alexandra C. Schmid for providing source codes to calculate coverage, sharpness, and contrast metrics from a specular image and Wiebke Siedentop and Annika Zentel for assisting with data collection. TM is supported by a Sir Henry Wellcome Postdoctoral Fellowship from Wellcome Trust (218657/Z/19/Z) and a Junior Research Fellowship from Pembroke College, University of Oxford. This research was also supported by “The Adaptive Mind”, funded by the Excellence Program of the Hessian Ministry of Higher Education, Science, Research, by the DFG SFB-TRR-135 “Cardinal Mechanisms of Perception” (project No. 222641018, TPs C1 and C2), and by a Marsden Fast Start Grant to KS from the Royal Society of New Zealand (project MFP-UOA2109). For the purpose of open access, the author has applied a CC BY public copyright license to any Author Accepted Manuscript version arising from this submission.

## Data access

The raw experimental data and source codes to analyze the data and reproduce figures will be available in a data repository at the time of publication.

